# EEG Resting-State and Event-Related Potentials as Markers of Learning Success in Older Adults Following Second Language Training: A Pilot Study

**DOI:** 10.1101/2020.10.03.324640

**Authors:** Maria Kliesch, Nathalie Giroud, Martin Meyer

## Abstract

In this pilot study, we evaluated the use of electrophysiological measures at rest as paradigm-independent predictors of L2 development for the first time in older adult learners. We then assessed EEG correlates of the learning outcome in a language-switching paradigm after the training, which to date has only been done in younger adults and at intermediate to advanced L2 proficiency.

Ten (Swiss) German-speaking adults between 65-74 years of age participated in an intensive three-weeks English training for beginners. A resting-state EEG was recorded before the training to predict the ensuing L2 development (Experiment 1). A language-switching ERP experiment was conducted after the training to assess the learning outcome (Experiment 2).

All participants improved their L2 skills but differed noticeably in their individual development. Experiment 1 showed that beta1 oscillations at rest (13-14.5Hz) predicted these individual differences. We interpret resting-state beta1 oscillations as correlates of attentional capacities and semantic working memory that facilitate the extraction and processing of novel forms and meanings from the L2 input. In Experiment 2, we found that language-switching from the L2 into the native language (L1) elicited an N400 component, which was reduced in the more advanced learners. Thus, for learners beginning the acquisition of an L2 in third age, language switching appears to become less effortful with increasing proficiency, suggesting that the lexicons of the L1 and L2 become more closely linked. In sum, our findings indicate that individual differences in L2 development and proficiency in older adults operate through similar electrophysiological mechanisms as those observed in younger adults.

## 1. Introduction

While many resources have been dedicated to establishing the now commonly accepted view that the learning of foreign languages is desirable for children and younger adults, little effort has been dedicated towards exploring the potential of this learning challenge for older adults (Pfenninger & Singleton, 2019). While the reasons for older adults to learn a new language are manifold, caution needs to be taken when applying research findings from younger adults to older learners. It has repeatedly been shown that L2 learning differs between younger and older adults, not only regarding the learning of new words (van der Hoeven & de Bot 2012), but also concerning the type of context required for successful word retention (Whiting, Chenery & Copland, 2011), the preference of implicit over explicit learning conditions, detrimental effects of either vision or hearing etc. (Cox, 2014). Learning a new language is one of the most complex cognitive tasks humans are able to perform, so that differences in L2 learning success are also likely to be a result of structural and functional brain changes commonly observed in healthy aging. For one, aging is accompanied by substantial brain atrophy, and it has also been associated with reduced capacities in a large number of cognitive skills, including but not limited to attention, executive functions, memory, problem solving and processing speed (Fijell & Walhovd, 2010; Salthouse 2012; Salthouse 2010). While there is an overall tendency of cognitive decline even in healthy aging, however, individual differences in cognitive performance increase with age and are also found in late L2 acquisition (Ingvalson, Nowicki, Zong, & Wong 2017; Kliesch et al., 2018; Mackey & Sachs 2012). In order to cater for the individual needs of older adults with varying L2 learning capacities, the first step is to understand the relationship between the aging brain and L2 development in third age.

Since understanding the between neural patterns and behavior can help us understand individual differences in cognitive abilities, L2 learning and age, neuroscientific research is an ideal complement in order to evaluate linguistic theories on late L2 acquisition. Despite this, the neurological substrate for these learner differences, particularly in older learners, is still largely unknown (Biedroń 2015). A useful tool to remedy this issue has proven to be the electroencephalogram (EEG). Its excellent temporal resolution of which allows us to record neural oscillations in the brain at rest and makes it possible to track changes in time-locked electrical brain activity associated with the processing of a second language. Of the two, investigating neural oscillations at rest is a promising way of approaching L2 development aptitude, that is, an individual’s sensitivity to L2 input that explains why some learners find it easier to acquire a new language than others, independently of motivational factors (Dörnyei & Skehan, 2002). Neural endogenous oscillations reflect stable aspects of the functional architecture that also underlie evoked oscillatory patterns, and have therefore been identified as an electrophysiological predictor of behavior (Campbell & Schacter, 2016). As with cognition, however, neural oscillations, too, are affected by age-related changes, so that the role of EEG indices can be particularly informative in older learners. On average, older adults show slower alpha activity (8-13Hz), lower amplitudes in the alpha and beta (14-30Hz) band and an increase in slower oscillations, that is, in the delta (1-4Hz) and theta (4-8Hz) range (Ishii et al., 2017). Reductions of beta power at rest have been related to alertness deficits and have been identified as markers of dementia progression (Coben, Danziger, & Storandt, 1985; Ishii et al., 2017), while overall indices of spontaneous electrophysiological activity have been found to be a reliable predictor of cognitive impairment and precursors thereof (Babiloni et al., 2011). At the same time, neural oscillations can be modified through cognitive training even in third age (Klados, Styliadis, Frantzidis, Paraskevopoulos, & Bamidis, 2016; Reis et al., 2016; Styliadis, Kartsidis, Paraskevopoulos, Ioannides, & Bamidis, 2015), and can therefore manifest great interindividual variability (Fernández et al., 2012). As a consequence, it is to be expected that changes in EEG resting indices are also likely to affect learning abilities in old adulthood. Despite these findings, there are – to the best of our knowledge – very few studies that have investigated EEG oscillations in the context of *learning* a new language, let alone in older learners who begin the L2 acquisition when they have already reached the third age.

In three EEG studies, neural oscillations at rest have been used as a predictor of L2 learning in younger adults, but there are no comparable studies with older learners. The first two of the three studies (Prat et al. 2016, 2018) used EEG indices (i.e. power in different frequency bands) to predict success of L2 learning in young adulthood (18-31yrs). In Prat (2016), the 16 participants were monolingual English speakers who completed an 8-week French course consisting of 16 30-minute sessions. L2 proficiency was assessed by recording the level each participant had reached at the end of the 16 sessions. Before the training, 5 minutes eyes-closed resting-state EEG data were collected, and power values were calculated across theta (4-7.5Hz), alpha (8-12.5Hz), beta1 (13-14.5Hz), beta2 (15-17.5), beta3 (18-29.5Hz) and low-gamma (30-40Hz) bands. Correlations between power values at each electrode and final L2 level showed that power values in the gamma, beta and theta bands, were predictive of subsequent individual differences in L2 learning. The beta1 band was the most predictive frequency band and yielded positive correlations between *r =* .60-.77. However, no correlations were found between EEG indices and changes in cognitive capacities. Prat et al.’s results are in line with those of Küssner et al. (2016), who found that beta power at various electrode sites predicted word recall in a foreign-vocabulary learning task, and did so at three different testing sessions. These studies indicate that resting-state EEG indices are a promising candidate for an electrophysiological measure of the above-mentioned L2 aptitude, since they provide a paradigm-free measure that predicts L2 development *before* the actual training. The current pilot study assesses whether in older adults, where heterogeneity in resting-state indices has been found to increase as a function of cognitive demands, the above findings can be replicated. The findings thereof help us understand whether L2 learning is qualitatively similar between older and younger adults from an electrophysiological point of view, which in turn can inform future research designs in terms of individualizability and customizability of training methods and materials for this age group.

In addition to understanding how the language acquisition progresses in older L2 novices, it is equally informative to investigate how the new language is integrated with the existing one in older learners at a given stage of L2 proficiency. Again, a previous study with younger adults could show that heterogeneity in L2 proficiency *after* L2 training manifests itself through differential processing of language switching between the newly learned and the native language. By means of electrophysiological correlates of L2 learning in younger adults, Van der Meij et al. (2011) could show that the electrophysiological response towards a language switch was indicative of the individual L2 level. The authors tested two groups of young adult monolingual Spanish EFL-learners, who, based on self-rating, qualified as intermediate or advanced L2 learners. Participants were visually presented with English (L2) sentences of the type *The house that we rented was <u>furnished</u> and felt <u>cozy</u>*, in which one of the adjectives could occur in L2 or L1 (Spanish). ERPs time-locked to the onset presentation of the critical adjective showed a clear N400 effect and a late-positive component (LPC) towards language switch in both proficiency groups. However, for the more proficient group, the N400 amplitude was larger and the effect showed a more frontal distribution. The authors interpreted their findings within the framework of the Revised Hierarchical Model (Kroll & Stewart 1994), which postulates that, in the early stages of L2 learning, there is a weak link between the L2 lexicon and the conceptual level but a strong one between L2 and L1 lexicons. Since the performed language switch was purely lexical (i.e. no semantic inconsistency was performed), the authors concluded that the less proficient learners manifested a stronger link between L1 and L2 words, thus facilitating language switching, while the proficient learners showed a pattern closer to that of balanced bilinguals. The participants in Van der Meij’s study, however, were younger adults, and it is possible that lifelong monolingualism in conjunction with decreased cognitive capacities leads to a more rigid language system in old age, which in turn could lead to reduced switching effects into the L1 or even the absence thereof, particularly at initial stages of L2 learning.

Thus, it remains to be tested, as we have done in the current pilot study, whether older adults would also display similar effects of language switching into their L1 and whether they would do so at very basic levels of L2 proficiency. These findings in turn may help L2 instructors and researchers select the primary teaching language and the appropriate amount of language switching in classrooms of older L2 learners, and help understand language-mixing errors or word retention difficulties in this age group.

## 2. Experiment 1

The aim of Experiment 1 was to replicate the studies by Prat et al., (2016, 2018) in older adults, and thereby examine whether pre-training resting-state EEG markers can predict L2 aptitude in third age learners following three weeks of L2 instruction only. Using EEG indices measured before the L2 course, we hypothesized – in line with Prat et al. (2016) – that power in the beta1 band in particular would predict individual L2 development.

### 2.1. Materials and Methods

#### 2.1.1. Participants

For this longitudinal study, we recruited ten healthy older participants (range = 65-73 yrs, *M* = 68.2 yrs, *SD* = 2.44, 4 women), all of whom were (Swiss-) German speakers, with no more than school knowledge of any language other than (Swiss-) German, little to no exposure to English, and who had not resided for more than three weeks in an English-speaking country during the past 40 years. None of them reported any history of present or past neurological, psychiatric, or neuropsychological disorders, and we excluded participants with hearing thresholds above 40dB on the better-hearing ear for frequencies lower than 500Hz, as this is the threshold considered to be disabling by the WHO. The study was approved by the local ethics committee of the University of Zurich. All individuals gave their written informed consent, and were refunded the course fee for their participation in the study.

Subsequently, all participants completed a 3-week English course for beginners, comprising a total of 60 hours distributed over all consecutive workdays within the course period (*d* = 15). Participants were trained in all four of the essential skills for L2 learning (i.e., speaking, writing, listening and reading) by a qualified English-as-a-foreign-language (EFL) teacher. One lesson per day was reserved for self-study and no additional homework was required. The structure and content of the training corresponded to that of a regular L2 course learners would receive at a language school for adults. Since the training was designed as an intensive course, the instructor had to be able to attend each learner individually so as to avoid some learners being outpaced by the others half way through the course, which would have rendered the course unteachable. Thus, for logistic reasons, and because redoing the course with a new set of participants would have added uncountable confounding factors (e.g. between-participants and teacher-participants group dynamics), we only report findings on one experimental group.

In the week before and again in the week after the training, resting-state EEG data were obtained from each participant and L2 proficiency was assessed via three language tests. In the week following the L2 training, an ERP experiment was carried out to investigate the N400 and LPC components as a response towards language switch and semantic incongruence, following the work of Van der Meij et al. (2011).

#### 2.1.2. Language Development

*To* reliably gauge L2 proficiency of English before and after the training based on language comprehension and production on the lexical as well as grammatical levels, learners completed three different L2 tests before and after the training. These will be described in the following.

##### Integrative L2 Knowledge

The C-Test, a language assessment, screening and examination tool, measures integrative L2 production skills; that is, a learner’s ability to infer missing information in a text where the natural information redundancy is reduced. For our study, the test consisted of five short, random, written texts in L2, the degree of difficulty of which was adjusted to the target level of the training (approximately basic level A2 as per CEFR (Council of Europe 2001)). In each of the texts, the second half of every second word was removed (see e.g., Aguado, Grotjahn & Schlak 2007; Coleman, Grotjahn & Raatz 2002), creating a total of 125 gaps that the participants had to fill in. Percentiles were calculated based on the number of correctly filled gaps.

##### Language Assessment Test Based on Course Book

The language training followed the text/workbook *Next A1* by Hueber Verlag regarding learning content, structure and exercises. The course book provides an online assessment test to measure vocabulary and grammar as well as basic listening comprehension with a target level of A1+ (CEFR). Participants completed the test in the lab with the presence of the experimenter ensuring that there were no problems based on computer illiteracy. Percentiles for each individual were calculated based on raw scores.

##### Listening Comprehension

To complement the listening tasks of the course book’s online assessment, which measures comprehension for letters, digits and gist only, a listening task was added to test comprehension on word and sentence levels. The test comprised twelve sentences to be translated by participants, each sentence corresponding to the level of difficulty of one unit in the book. Two points were awarded for sentences translated correctly in both content and form, and percentiles were calculated for each individual.

##### Corrected L2 Development

Given the omnipresence of English loan words in (Swiss-) German or in advertisements and the media in general, a certain degree of previous L2 skill was practically unavoidable. In order to control for the differences in previous L2 proficiency, the language tests were carried out both before and after the training. A principal component analysis revealed that all L2 scores loaded on the same factor, which explained 82% of the variance within the three language tests. Therefore, L2 proficiency was calculated as the mean over all language tests. L2 development was not calculated as the difference between post- and pre-scores, since this would have rewarded learners with low initial L2 skills. Instead, because L2 improvement becomes more difficult and thus decelerates with increasing proficiency, it was corrected to reflect the percentage of maximum attainable improvement for each learner, as follows:

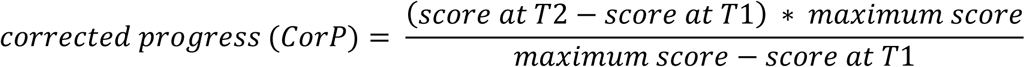

Thereby, a learner starting with previous L2 knowledge of 30 points who improves by 20 points has a higher CorP (CorP = 28.57) than a learner with zero previous knowledge who also improves by 20 points (CorP = 20.00).

#### 2.1.3. Resting-State EEG Recording

Before and after the training period, 8 minutes of alternating 2 minutes eyes-open/2 minutes eyes-closed resting-state EEG data were collected from participants, using 128 Ag/AgCl electrodes embedded in an elastic cap (Electro-Cap International Inc. Eaton, OH, USA), and recorded with the high resolution BioSemi ActiveTwo EEG system (BioSemi B.V., Amsterdam, Netherlands). Each participant was seated in a soundproof, electrically shielded room, approximately 80cm away from the computer screen. The electrical brain activity was recorded with a sampling rate of 512Hz. Impedances were generally kept below 20kΩ and the signal was filtered online with a band pass filter of 0.1-100Hz. Preprocessing was conducted using the BrainVision Analyzer 2.1 (Brain Products, http://www.brainproducts.com). A low cutoff filter of 0.1Hz (12dB) and a high cutoff of 30Hz (48dB) were applied offline. For the extraction of power means per frequency band, we followed the procedures described in Prat et al. (2016), again using BrainVision Analyzer. Accordingly, we only used the eyes-closed resting-state segments (a total of 4 minutes) from the 8-minute recording to allow direct comparability of results with Prat et al. (2016). The data were then segmented into 2-second-epochs while automatically skipping bad intervals. To correct blinks and saccades, an independent component analysis (ICA) was applied (Jung et al. 2000), and semi-automatic artifact rejection within each channel was used to eliminate noisy segments. Data were rejected from 200ms before to 200ms after a given event if the gradient exceeded a voltage step of 50μV, if the maximal difference of values in a 200ms interval exceeded 200μV, and if activity in 100ms intervals was not lower than 0.5μV. Channels that would have caused more than 10% of an individual’s data to be rejected due to artifacts were replaced via topographic interpolation; that is, their activity was simulated by averaging the activity of the adjacent electrodes. Finally, all activity was re-referenced to the average reference.

#### 2.1.4. EEG Analysis

Using the Fast Fourier Transform, we extracted the mean power in μV^2^ across five different frequency bands by averaging the resulting power spectra across all epochs. Following Prat et al. (2016), power means were calculated separately for theta (4-7.5Hz), alpha (8-12.5Hz), beta1 (13-14.5Hz), beta2 (15-17.5Hz) and beta3 (18-29.5Hz) for each frequency pool and in each participant. Following Prat et al. (2016) and Doppelmayr et al. (2002), all power means were log-transformed and used as such in all further analyses.

For the statistical analysis of the data, power means per subject, electrode pool and frequency band were subjected to a multilevel model (Bliese 2009, Bryk & Raudenbush 1992), as they do not require normal or parametric data, and allow controlling intraclass correlation. Intraclass correlation coefficient (ICC) was used as a measure of whether measurements of ERP amplitude were independent within subjects in order to determine the necessity of multilevel (i.e. mixed) models. Power values were nonindependent in subjects, ICC(1) = .37, *F*(9.290) = 18.76, *p* < .001, which according to Cichetti provides fair significance that measurements were nonindependent within subjects (1994). In the Mixed Model, resting-state oscillatory power was used as independent variable, predicted by the main factors Frequency Band and L2 Development, with random effects for subjects. All values were standardized before entering into the models. This method allowed us to account for intraclass correlation within subjects and to include all frequency bands within the same model, thereby reducing type I error based on multiple comparisons. The alpha band was used as reference frequency in the model, since we did not expect any significant correlation between alpha and L2 development (Prat et al. 2016, 2018). The model did not differentiate between electrodes in order to avoid overfitting and to determine the overall relationship of L2 learning and EEG indices, independent of electrode position. Wherever possible, and as recommended by the American Statistical Association (ASA), we report confidence intervals instead of *p*-values, given the well-known shortcomings of the latter in providing a good measure of evidence for a model or hypothesis, particularly because it is confounded by the number of observations in our sample, which would likely result in a type II error (Ranstam 2012; Wasserstein & Lazar 2016). Furthermore, *p*-values have repeatedly been shown to be problematic if not altogether unnecessary in mixed models (Bates 2014). In contrast, confidence intervals allow us to make statements as to which effects are likely to exist in the population and provide a good measure of how precise the sample statistic is. For all statistical analyses we used the program R (http://www.r-project.org).

### 2.2. Results

#### 2.2.1. Individual Differences in Language Learning

Consistent with previous research, even though all participants increased their L2 skills over the course of the 3-week training, there was large individual variability in the L2 improvement, ranging from 16.66% to 91.86% of the maximum attainable improvement (*M* = 45.33, *SD* = 24.92). The overall L2 development from T1(*M* = 41.09, *SE* = 6.96) to T2(*M* = 65.04, *SE* = 7.30) was significant (*t*(9) = 6.33, 95% CI [17.01, Inf], *r* = .9). Individual L2 development is depicted in Figure 1. The relationship between initial L2 knowledge (which were very basic in all participants) and the degree of L2 development over the course was non-significant (*r* = 0.56, 95% CI [−0.11, 0.88]). Indeed, some participants who started with lower L2 skills at T1, after the training had surpassed other learners who began with higher levels of L2 knowledge. In the following, we assess whether the same learner differences are manifested in the EEG experiments.

**Fig. 1.**
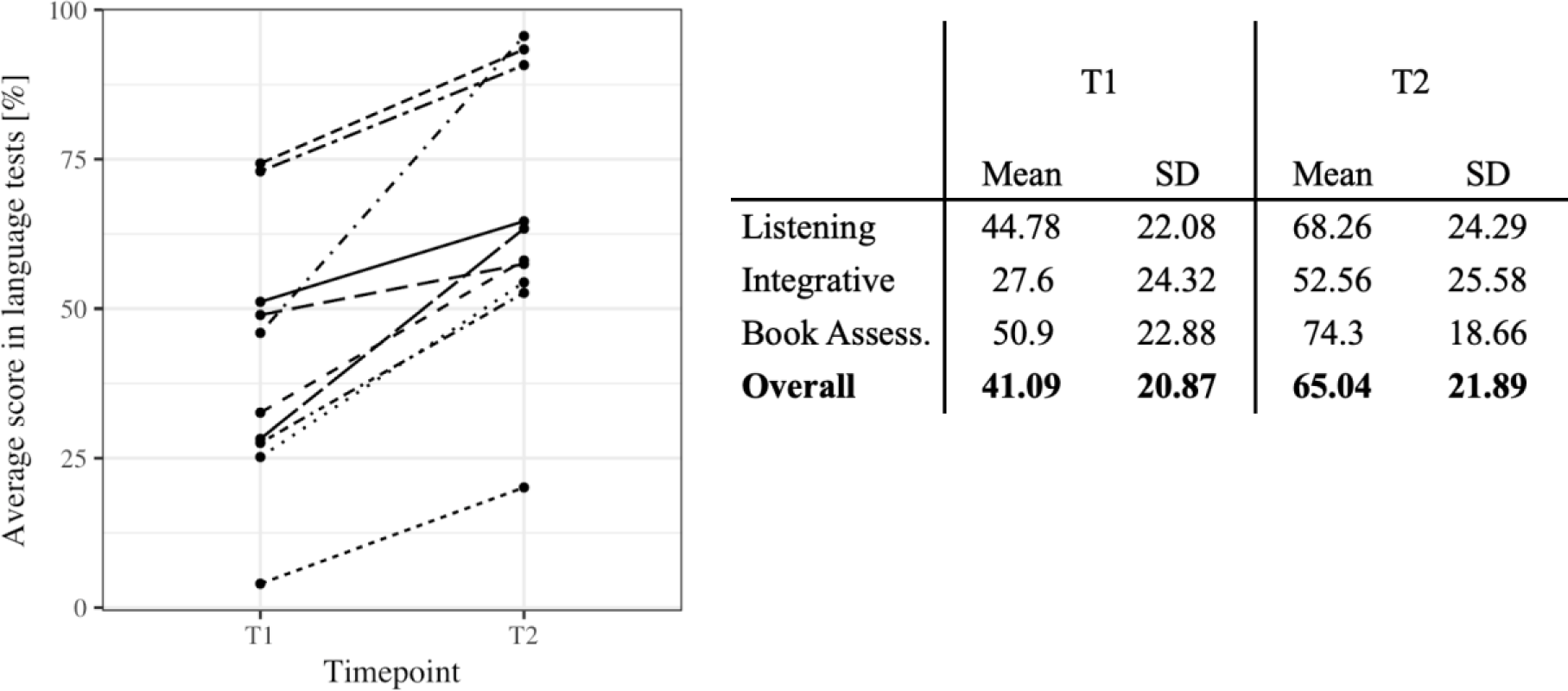
Left: Individual language scores before (T1) and after the training (T2). Each line represents change in L2 proficiency of a given learner. Right: Overview of individual L2 scores at T1 and T2 and the Corrected L2 Development.

#### 2.2.2. Resting-state EEG indices

Table 1 shows the results of the multilevel model used to calculate the main effect of L2 development and frequency as well as their interaction on the resting-state EEG power. The resting-state data used were those before the training, so as to assess whether EEG indices would predict the L2 development. Consistent with Prat et al. (2016, 2018), we found a positive relationship between language learning and all assessed frequency bands, which explained 33% of the variance observed. However, only in the beta1 band, power was significantly predicted by L2 development. This finding is in line with Prat et al. (2016), who found the highest correlation coefficients between L2 learning rate and EEG indices in the beta1 band (*r =* .60-.77) and Prat, Yamasiki and Peterson (2018) who found a correlation of L2 learning rate with pre-training power in the beta band over right posterior electrodes (*r*_*s*_ = .39). Following the findings by Prat et al. (2016) and taking their analysis a step further, in addition to investigating EEG measures as predictors, we also investigated power changes in the beta1 band via a paired *t*-test of beta1 power in pre and posttests. The difference between pre- and post-values in beta1 power was correlated with the L2 development in order to assess the relationship between stability in EEG indices and L2 outcome.

**Table 1:**
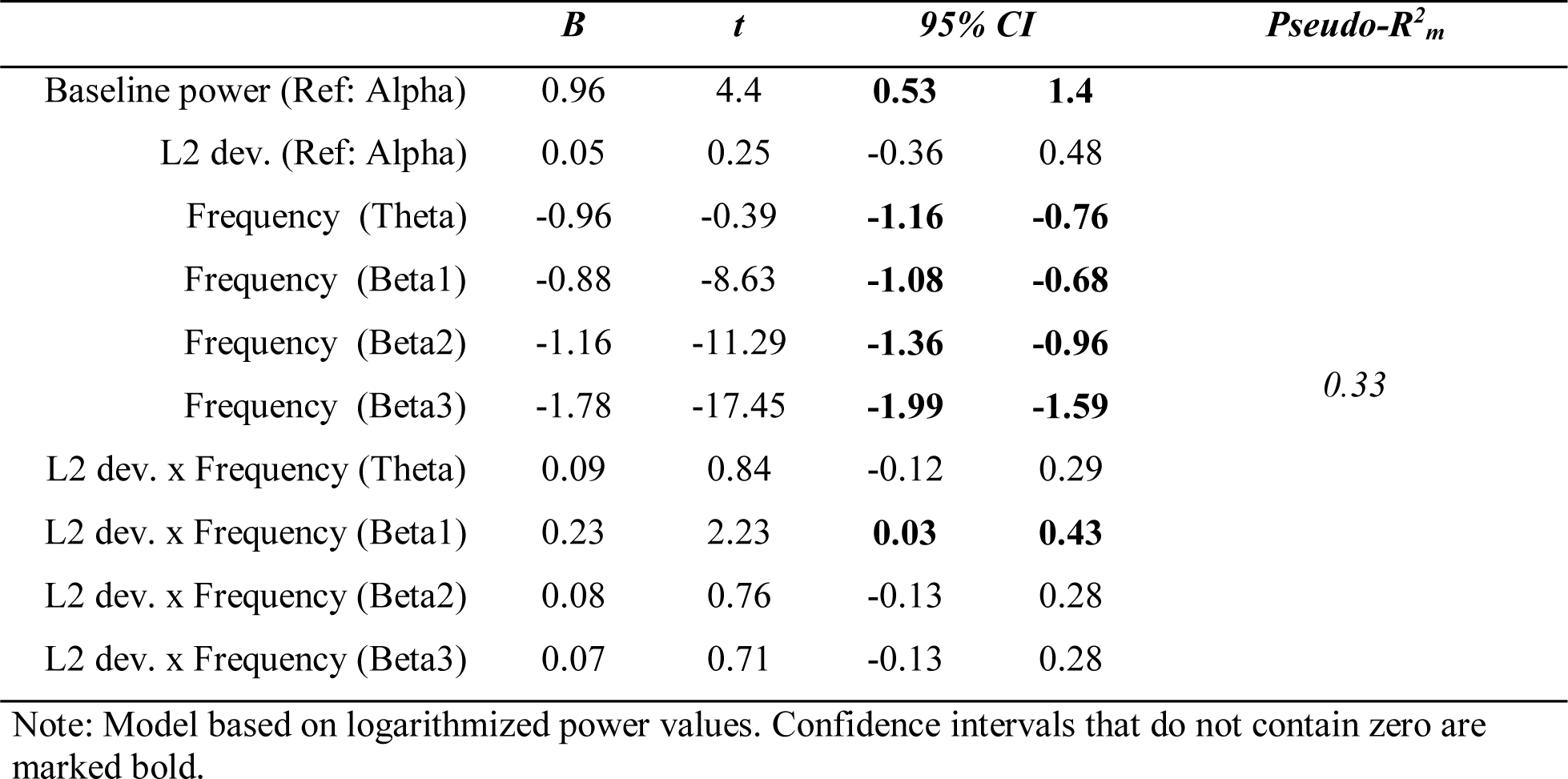
Result from Multilevel Model predicting L2 development based on averaged power in each of the frequency bands.

|  | *B* | *t* | *95% CI* |  | *Pseudo-R<sup>2</sup><sub>m</sub>* |
| --- | --- | --- | --- | --- | --- |
| Baseline power (Ref: Alpha) | 0.96 | 4.4 | **0.53** | **1.4** | 0.33 |
| L2 dev. (Ref: Alpha) | 0.05 | 0.25 | -0.36 | 0.48 |  |
| Frequency (Theta) | -0.96 | -0.39 | **-1.16** | **-0.76** |  |
| Frequency (Beta1) | -0.88 | -8.63 | **-1.08** | **-0.68** |  |
| Frequency (Beta2) | -1.16 | -11.29 | **-1.36** | **-0.96** |  |
| Frequency (Beta3) | -1.78 | -17.45 | **-1.99** | **-1.59** |  |
| L2 dev. x Frequency (Theta) | 0.09 | 0.84 | -0.12 | 0.29 |  |
| L2 dev. x Frequency (Beta1) | 0.23 | 2.23 | **0.03** | **0.43** |  |
| L2 dev. x Frequency (Beta2) | 0.08 | 0.76 | -0.13 | 0.28 |  |
| L2 dev. x Frequency (Beta3) | 0.07 | 0.71 | -0.13 | 0.28 |  |

Note: Model based on logarithmized power values. Confidence intervals that do not contain zero are marked bold.

**Table 1:** Fixed effect estimates for the best-fitting model for the interaction of L2 development and pre-training qEEG power over all frequency bands
|  | *B* | *t* | *95% CI* |  | *Pseudo-R<sub>m</sub><sup>2</sup>* |
| --- | --- | --- | --- | --- | --- |
| Baseline power (Ref: Alpha) | 0.96 | 4.4 | **0.53** | **1.4** | 0.33 |
| L2 dev. (Ref: Alpha) | 0.05 | 0.25 | -0.36 | 0.48 |  |
| Frequency (Theta) | -0.96 | -0.39 | **-1.16** | **-0.76** |  |
| Frequency (Beta1) | -0.88 | -8.63 | **-1.08** | **-0.68** |  |
| Frequency (Beta2) | -1.16 | -11.29 | **-1.36** | **-0.96** |  |
| Frequency (Beta3) | -1.78 | -17.45 | **-1.99** | **-1.59** |  |
| L2 dev. x Frequency (Theta) | 0.09 | 0.84 | -0.12 | 0.29 |  |
| L2 dev. x Frequency (Beta1) | 0.23 | 2.23 | **0.03** | **0.43** |  |
| L2 dev. x Frequency (Beta2) | 0.08 | 0.76 | -0.13 | 0.28 |  |
| L2 dev. x Frequency (Beta3) | 0.07 | 0.71 | -0.13 | 0.28 |  |

Note: Model based on logarithmized power values. Confidence intervals that do not contain zero are marked bold.

When comparing beta1 values before and after the language course, the paired *t-*test revealed a negative trend; that is, a decrease in beta1 power between T1 (*M* = −0.85, *SE* = 0.35) and T2 (*M* = −0.89, *SE* = 0.32), (*t*(59) = −1.91, 95% CI [−0.08, 0.00], *r =* .24). However, there was no correlation between the change in beta1 power over the course and eventual L2 attainment (*B* = 0.05, 95% CI [−0.21, 0.32], *t*(58) = 0.41).

### 2.3. Discussion of Experiment 1

In this experiment, we investigated whether brain activity at rest predicts individual L2 aptitude in older adults, who only begin to learn an L2 when they have already reached the third age. Our results demonstrated that endogenous brain activity, as measured by beta1-power in the resting-state EEG before the training, was a reliable predictor of individual L2 progress, since participants with higher power in the beta1 band prior to the training also showed larger L2 development during and after the training. These findings are consistent with those of Prat et al. (2016), who also found correlations to be strongest between L2 learning rate and beta1 power. In their study, correlations between L2 learning rate and power in the beta2 band were smaller but still significant, while those in the beta3 were not significant after FDR correction. In a similar publication from 2018, Prat, Yamasaki and Peterson reported findings from the same study but using the entire beta band for correlations with learning rate, total speech attempts and speaking accuracy. Unsurprisingly, since the beta1 band forms part of the larger beta band, positive correlations were also found for overall beta and L2 measures. Similarly, our findings are in line with Küssner et al. (2016), who found baseline resting-state beta (14-35Hz) power to be a predictor of word recall in a foreign vocabulary task, and are compatible with those of Kepinska et al. (2017), who found that learning of an artificial grammar was more successful in learners who showed higher functional connectivity in the beta band (13-29Hz) during the learning phase.

Since our results and that of Prat et al. (2016) suggest a differential role for beta1 oscillations in L2 learning as opposed to beta2 or beta3 oscillations– both in younger and older adults – we will first address those previous studies that report on the beta1 band in particular and its relationship with cognitive functioning. Egner & Gruzelier (2004) could show that, in younger adults, neurofeedback training of low beta frequencies (12-15Hz and 15-18Hz) led to increased perceptual sensitivity, reduced omission errors and faster reaction times than in a non-neurofeedback control group. Similarly, Egner & Gruzelier (2001) found that training these same frequency bands also lead to increases in P300 ERPs, a component which is heavily influenced by attention (Szuromi, 2011). In line with these findings, Vernon et al. (2003) reported improved accuracy in focused attentional processing and improved performance in a semantic working memory task following beta1 training (12-15Hz). In line with these studies, Gruzelier et al. (2014) in their review associate power in the 12-15Hz band (referred to as sensori-motor-rhythm) with attentiveness, sustained attention, semantic working memory, declarative memory and reduced hyperactivity.

Even though the beta1 band appears to have a distinctive role in cognitive performance and L2 acquisition, few studies actually discuss the absence of effects in the upper beta bands. Park et al. (2008), for instance, found that power in all beta bands was reduced in patients suffering from Alzheimer’s disease (AD), but also found that the effect was strongest in the beta1 band, smaller in the beta2 band, and least pronounced – but still significant – in the beta3 band. The authors, however, do not discuss this graded effect. Accordingly, Hogan et al. (2003) report an increase of beta1 power in both healthy older adults and adults suffering from AD during memorization phases as a function of working memory load, but only AD patients also increased beta2 and beta3 power. Again, the authors omit any discussion of these differences. Finally, the study by Lindau et al. (2003) also found distinctive patterns of oscillatory power decrease in the beta1-beta3 bands between healthy older controls, patients suffering from AD and those suffering from frontotemporal dementia, but again, the authors fail to discuss those.

Therefore, it appears that to date, there is no theoretical model on the distinctive functions of beta1-beta3 oscillations in cognition and language processing, but the evidence appears to suggest that their function in cognition may vary. Since most studies, however, do not make this distinction, we will also discuss results on the overall beta band if their frequency range of interest included 13-14.5Hz (our beta1 band). Similar to beta1 studies, those on the overall beta band have linked beta power to attentional resources that naturally fluctuate as a function of attentional resources and mental fatigue throughout the day. For instance, Jap et al. (2009) and Liu, Zhan & Zheng (2010) showed that beta power decreased as a function of fatigue following exhausting tasks, such as prolonged monotonous driving or repeated cognitive tasks. Accordingly, Kepinska et al. (2017) found that artificial grammar learning was more successful in 1) participants who had high functional connectivity in the beta band (13-29Hz) during the learning phase, and 2) participants who had high beta power right at the beginning of the learning phase – and not increasing throughout the task. The authors associated beta with improved memory encoding during operations of high memory load. In line with these findings, Engel and Fries (2010) postulate that beta band activity relates to maintaining the current motor and cognitive set, which can be understood as signaling the status quo via endogenous top-down processes. Typical top-down processes that require maintaining of a cognitive set include working memory tasks and attention, both of which have been linked to beta band activity (Engel & Fries, 2010). In particular, a study by Gola et al. (2013) showed that older adults who manifested a decrease in beta power during the anticipatory period of a visual attention task were significantly poorer performers than those who showed a beta-band power increase. These findings led the authors to conclude that beta power is associated with activating and sustaining attentional processes and, most importantly, that this relationship is present both during tasks as well as at during rest. As Engel and Fries (2010) hypothesized, beta band activity should be particularly high during a resting state in which there is no expectation of ensuing change in the sensorimotor set. Our results can be reconciled with both this hypothesis as well as the findings of Gola et al. (2013): If indeed beta power at rest is indicative of an endogenous, permanently oscillating attentional state, our results would indicate that intrinsic attentive capacities are an essential prerequisite for successful L2 learning.

There is ample evidence to suggest that it is precisely selective attention that is one of the key skills enhanced in bilinguals as compared to monolinguals (Bialystok 2009), and that it constitutes a significant predictor for L2 acquisition in adulthood (Batterink & Neville 2014, Issa & Morgan-Short, 2018). L2 learners acquire a new language based on the input they read or hear, but some features of the input become output only at very late stages of L2 acquisition, and some forms even fail to become intake altogether. According to Ellis (2006), the reason for this is “learned attention” in the L2 acquisition, which retards or even prevents the noticing of fragile features of L2 due to factors intrinsic to languages, such as salience or cue competition. Features, such as the third person singular “-s” of English, which are redundant (*He eat* an apple* would still be comprehensible) are notoriously difficult to acquire, as they are non-salient and therefore require increased levels of selective attention in order to be detectable in the language stream. If selective attention in a given learner is low, only the most obvious cues in the input may be acquired, which in turn may be sufficient for everyday communicative survival (Ellis, 2006) and thus impede L2 progress. As a consequence, the slower learners of our study are likely to have suffered from reduced attentional resources, as reflected in the lower power values in the beta1 band. Thus, our findings are not only in line with Prat et al. (2016, 2019), but are also consistent with previous findings on the relationship between beta1 oscillations, attentional processing and semantic memory, confirming that beta-activity in the brain at rest can be used as a paradigm-free tool to predict L2 development.

## 3. Experiment 2

In the second experiment, we aimed to determine whether electrophysiological measurements already differentiate varying levels of L2 proficiency after L2 training of only three weeks and in older adults, thus providing insight into the question of how a newly learned language in old adulthood is processed in relation to the native language (L1), and thereby informing theories on how the new language is stored at initial stages of L2 learning. To this end, we conducted a visual language switching experiment after the training similar to that of Van der Meij et al. (2011). English (L2) sentence were presented, in which the target word was either in L2 or L1, and either semantically congruent or incongruent with the rest of the sentence. We aimed to test the following hypotheses: In line with Van der Meij et al. (2011), we hypothesized an N400 effect as well as an LPC (late-positive component) towards language switch. Our learners being L2 beginners, however, we expected to find an inverse relationship between language-switching effects and L2 proficiency, assuming that at a very basic L2 proficiency, the link between L1 and L2 lexicons does not yet exist but strengthens with increasing proficiency until resembling that of Van der Meij’s intermediate learners. In addition, we hypothesized that an effect of semantic incongruence would either be reflected by a monophasic LPC or a biphasic N400-LPC pattern, as these have been shown to be elicited by similar semantic violations (e.g. Bornkessel-Schlesewsky & Schlesewsky 2008). The participants, the language training and the language tests were the same as in Experiment 1.

### 3.1. Materials and Methods

#### 3.1.1. Measure of Language Proficiency

In contrast to Experiment 1, in Experiment 2 we applied a different approach to test L2 proficiency after the training. Hence, we did not use the *Corrected L2 Development Score* but the mean over all L2 tests after the training to gauge L2 proficiency rather than L2 development, as we were assessing the momentary processing of language switching as a learning outcome and not the longitudinal development of L2 processing.

#### 3.1.2. ERP Stimulus Material

For the stimuli, following Van Der Meij et al. (2011), a 2 (*switch* vs. *no-switch*) x 2 (*congruent* vs. *incongruent*) design was used, with a total of 320 English sentences of nine to 12 words (see Figure 2). Sentence structure was the same for all stimuli; that is, a compound sentence that included a subordinate relative clause (e.g., “The girl that does not talk much writes a book”). The last word of each sentence could either occur in English (*no-switch*, 80 sentences) or in German (*switch*, 80 sentences, e.g., “The girl that does not talk much writes a Buch”), and could be semantically congruent (*congruent*, 80 sentences) or incongruent with the rest of the sentence (*incongruent*, 80 sentences, e.g., “The girl that does not talk much writes a potato” / “The girl that does not talk much writes a Kartoffel”). Since word order is not identical in English and German, the code-switch was performed in the last part of the sentences, where word order is grammatically correct in both languages. Each sentence appeared in all four conditions, and in order to prevent participants from recognizing sentences from earlier iterations, the stimulus material was made up of no more than 20 nouns and 15 verbs in total, which were reassembled into 80 different sentences via an automatic algorithm that forced each verb to appear in two sentences and each noun to appear in four sentences, twice as subject and twice as object. All words that appeared in the stimuli were part of the course book curriculum, and the instructor ensured familiarity with the terms over the course of the training. For the sentence-final target word, only nouns that were orthographically different in at least two letters between English and German were included, and there were no “false friends” (e.g., Gast – guest, \**Handy* – handy). Average frequency of the words was not taken into consideration, since their use in the classroom environment is not representative of that of a native speaker of English or German, but it was ensured that all words were covered in the course curriculum.

**Fig. 2.**
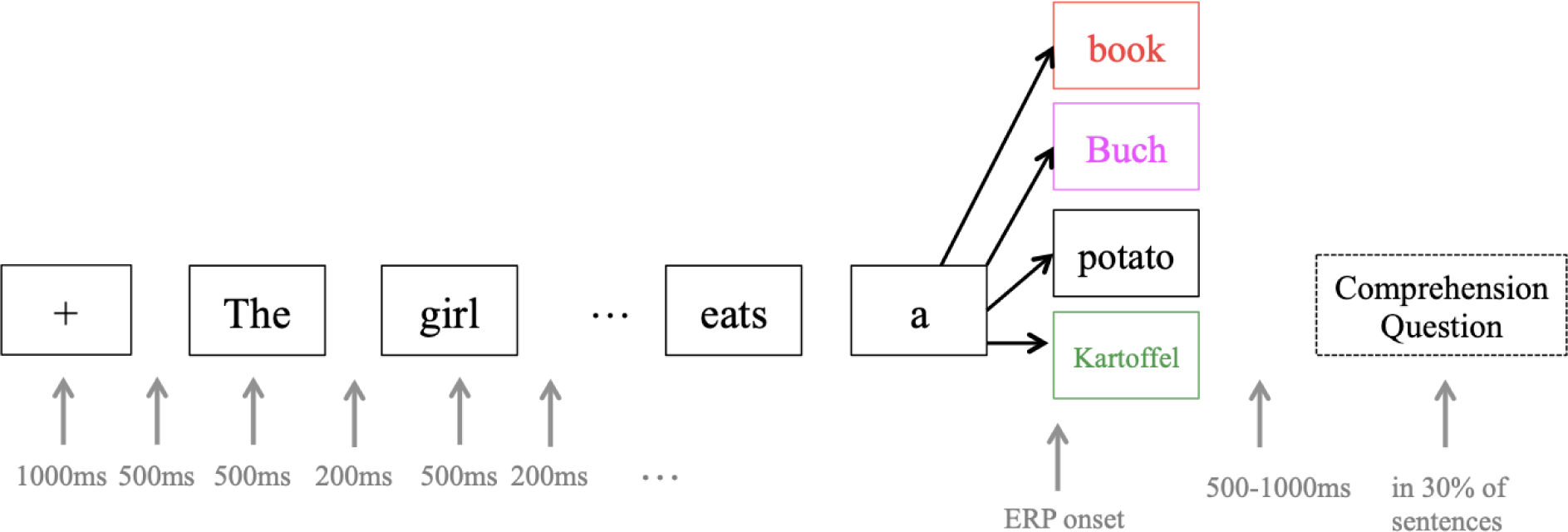
ERP experiment design. Each target word (e.g. book) appeared in at least four different sentences and conditions in order to avoid prediction effects. ERPs were recorded from the onset of the target word. The colors used for target words are the same as in Fig. 4: red = no-switching incongruent, pink = switching incongruent, black = no-switching congruent, green = switching congruent.

#### 3.1.3. EEG Recording and Preprocessing

EEG data were collected using the same system and settings as for the resting-state data. The presentation of stimuli was controlled via the Presentation software (Version 18.0, http://www.neurobs.com). Similar to the study of Van der Meij et al., (2011), sentences were presented visually one word at a time in a grey-green lowercase font against a black background. Each sentence was preceded by a “+” sign shown for 1000ms, followed by a blank screen of 500ms. Each word was shown for 500ms with a blank screen of 200ms between words. At the end of each sentence, a blank screen of a jittered duration of 500-1000ms was inserted to ensure onset asynchrony between sentences. Participants were instructed to read the sentences for content, and practiced the task in the presence of the experimenter. After 1/3 of the sentences, participants’ comprehension of the content was assessed with the question *Hat der letzte Satz Sinn gemacht?* (*Did the last sentence make sense?*), and answers had to be indicated via the left (“No”) or right (“Yes”) arrow key. The preprocessing procedure was the same as in Experiment 1.

#### 3.1.4. ERP Analysis

After preprocessing, the ERP data was segmented for each condition (*switch, no-switch, congruent, incongruent*) from 100ms before to 1000ms after the onset of the final noun and baseline corrected with regard to the 100ms pre-stimulus baseline. In order to be able to detect the expected N400 and LPC, two time windows of interest were determined through a mixed hypothesis- and data-driven approach (see Results). The extracted windows were 300-450ms for the N400 (Van der Meij et al. 2011) and 600-800ms for the LPC (Bornkessel-Schlesewsky et al. 2015; Dowens, Vergara, Barber, & Carreiras 2010). Since the effects were expected to be distributed over large areas of the scalp (Van der Meij et al. 2011), electrode pools were formed by subsuming electrodes of similar activity (see Results), which then served as regions of interest in all further analyses.

For the statistical analysis of the ERP data, the mean voltage amplitudes relative to the start of the critical noun were subjected to a multilevel model (Bliese 2009, Bryk & Raudenbush 1992). All values were standardized before entering into the models. Intraclass correlation for the time window between 300-450ms was *ICC* = .70, and *ICC* = .69 for the time window between 600-800ms, which according to Cichetti provides good significance that amplitudes were nonindependent within subjects (1994). In both time windows, the random intercept model fitted the data better than the random intercept-and-slope model, so that only random participant-intercepts were used. The model included main effects for Switch (switch, no-switch) and Congruency (congruent, incongruent) as well as their interaction as within-subject factors, and featured age as a control variable. In addition, a linear regression model was computed to assess the relationship between L2 proficiency after the training (T2) and the respective amplitude difference between switch and no-switch conditions, and between congruent and incongruent ones.

### 3.2. Results

#### 3.2.1. Accuracy

In the ERP experiment, participants’ accuracy in determining whether the target sentences made sense was 81.39%. All participants scored over 75% except for one learner, who only reached 57.8%. However, there was no significant difference in the model fit between the model using proficiency as a predictor and the one using accuracy for either of the ERP components (N400, LPC), as assessed via the Bayesian information criterion (BIC). In addition, there was a strong correlation between accuracy in the EEG-Experiment and L2 proficiency after the training (*r =* 0.86, 95% CI [0.58, 1.00]), so that we decided to retain all of the data in the mixed models calculated for each ERP component.

#### 3.2.2. Event-related potentials

Since the expected N400/LPC effects are known to be distributed over relatively large areas of the scalp (Van der Meij et al. 2011), we used a data-driven approach to define one electrode pool for each ERP component, combining adjacent electrodes that visibly reflected the expected activity. In accordance with Van der Meij et al. (2011), we expected to find a negativity around 400ms (N400) and a positivity around 700ms (LPC) as an effect of language switch, and postulated semantic incongruence to be reflected in either an N400 modulation or a late positivity around 700ms (Bornkessel-Schlesewsky & Schlesewsky 2008). Figure 3 shows topographic difference plots between conditions switch and no-switch, and between congruent and incongruent conditions, respectively. Based on these topoplots, a central electrode pool was formed to subsume all activity related to the N400 component, and a parietal pool was used to analyze LPC characteristics (see Figure 3).

**Fig. 3.**
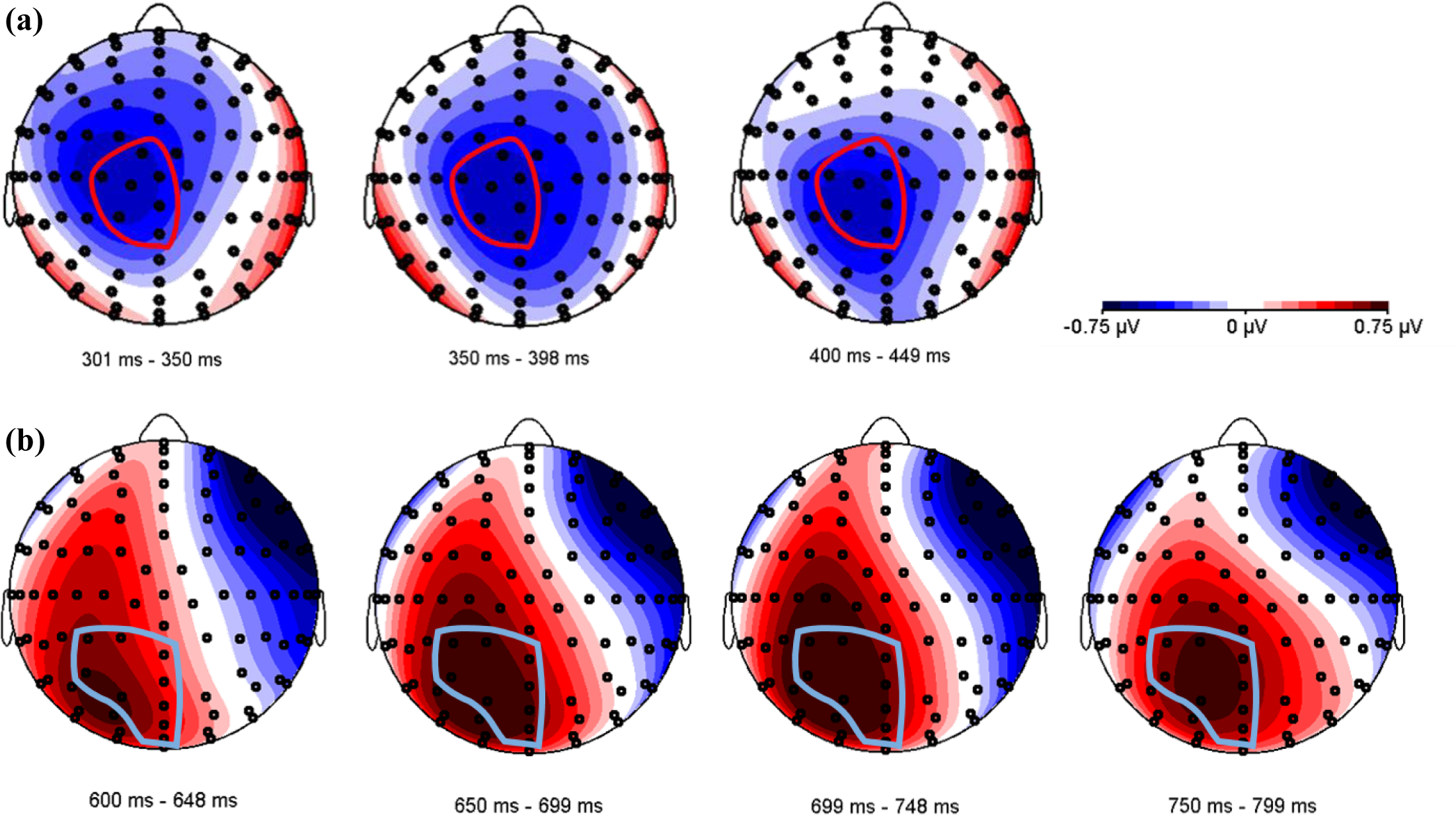
Topographical distribution of two ERP components. (a) N400 effect in the time window between 300-450ms. The voltage maps were obtained for the grand average values of language switch minus no-switch conditions. The electrode pool selected for further analyses is circled in red, and includes electrodes A1 (Cz), A2, A3 (CPz), D1, D14 (C1), D15 and D16. (b) LPC in the time window between 600-800ms. The voltage maps were obtained for the grand average values of incongruent minus congruent conditions. The electrode pool selected for further analyses is circled in light blue, and includes electrodes A3 (CPz), A4, A5, A6, A20, A21(POz), D16 (Fp2) and D17(FPz).

Figure 4 shows ERPs time-locked to the onset presentation of the final noun, averaged over all participants for the four experimental conditions (language switching, no language switching, semantic congruence, semantic incongruence), plotted in the two electrode pools.

**Fig. 4.**
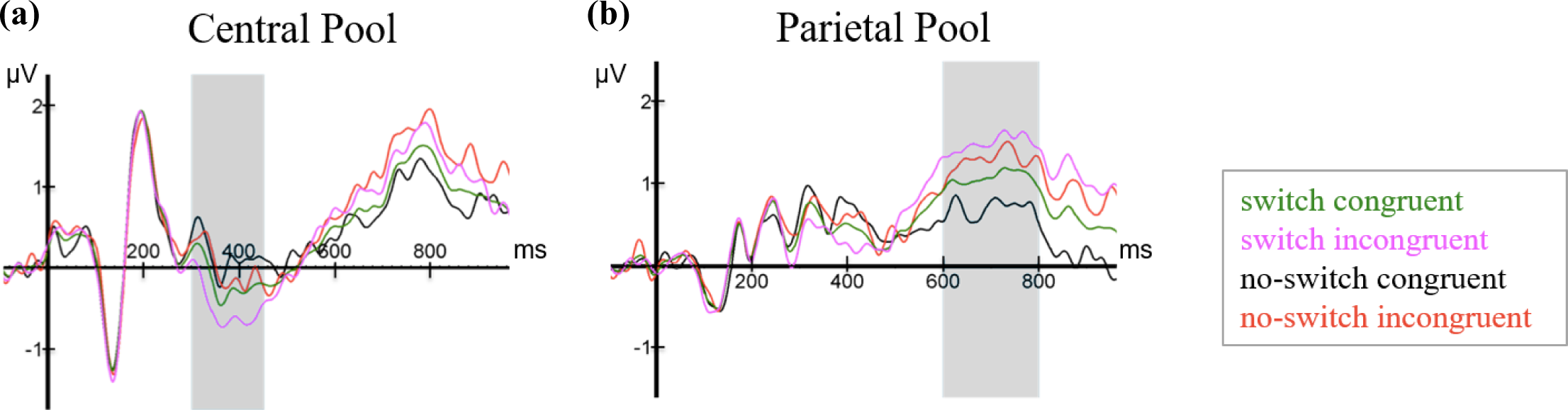
ERPs elicited by the last word in all four conditions (switch, no-switch, congruent, incongruent), plotted in the (a) Central and the (b) Parietal electrode pools.

At the Central electrode pool, the N400 and LPC components are clearly visible, although overall amplitudes are noticeably small compared to similar studies (Bornkessel-Schlesewsky et al. 2015; Van der Meij et al. 2011). The N400 is visibly scaled, such that average amplitudes were largest for the switch-incongruent condition, smaller for the switch-congruent condition, smaller still for the no-switch-incongruent condition and smallest for the no-switch-congruent condition. Thus, relative to no language switch, the two language switch conditions elicited a conspicuously larger negativity between 300-450ms after word onset, which is most prominent in the Central pool. In addition, already starting 500ms post-target word presentation, but more visibly after 600ms, a positivity at the Parietal pool with a duration of approximately 200ms and peaking around 700ms shows more positive values for semantically incongruent conditions than for congruent ones. Based on the visual inspection, the Central pool was chosen for all further analyses of the N400, and the Parietal pool for those of the LPC. As shown in Figure 5 and consistent with Van der Meij et al. (2011), the N400 effect occurs in the time window between 300-450ms. Therefore, we used this same time window for all further statistical analyses of the N400 component. For the LPC, we used a data-driven approach, as the incongruent condition did not form part of Van der Meij et al.’s study (2011). As can be appreciated from the ERPs, the LPC is clearly discernible from approximately 600ms onwards. We chose a time window of 200ms (600ms-800ms). This was deemed sufficient to detect the expected effect in each learner based on existing literature on the LPC (Bornkessel-Schlesewsky et al., 2015; Dowens et al., 2010).

**Fig. 5.**
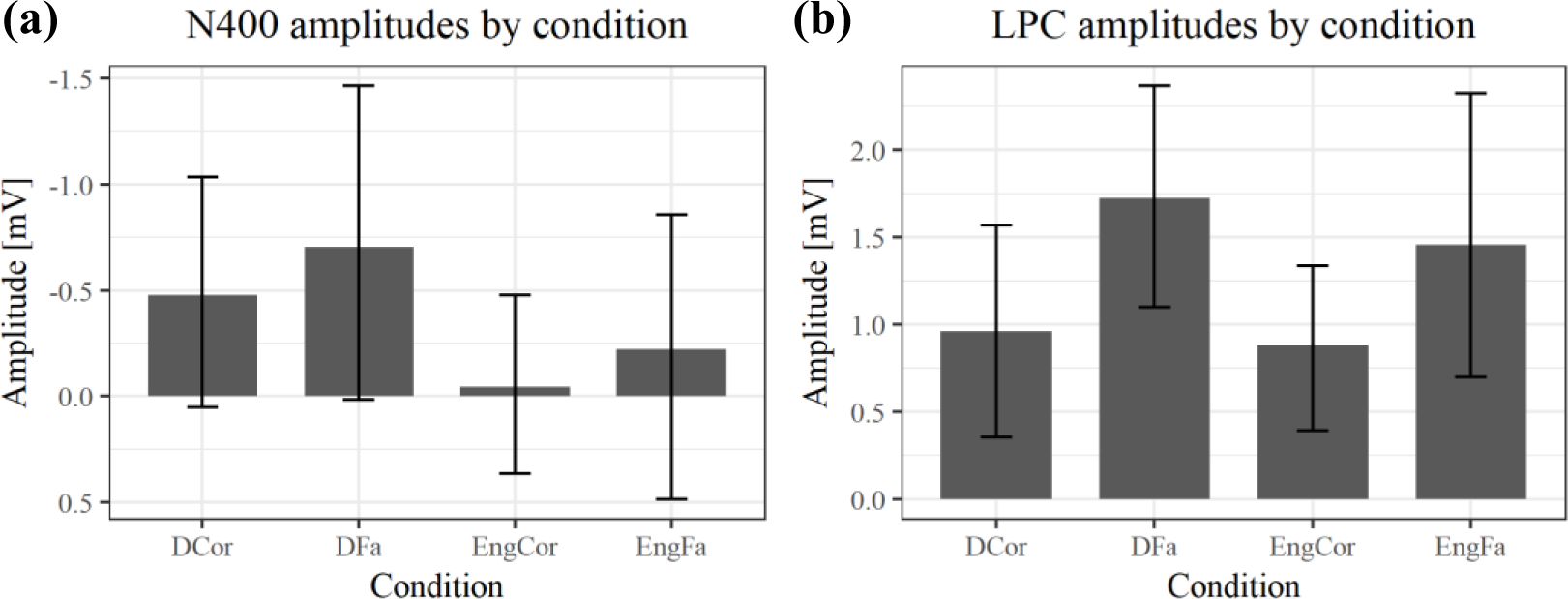
Bar charts of ERP amplitudes in each of the four experimental conditions averaged over all subjects. Plot (a) shows grand average amplitudes and their standard error for the time window between 300-450ms, Plot (b) those in the time window of 600-800ms. For the N400, there is a significant difference between language conditions, that is, between no switching (*Eng*) and switching (*D*), while in the LPC component, the significant difference was between congruent (*Cor*) and incongruent (*Fa*) conditions. As the error bars show, there appears to be a large overlap between conditions in the N400 time window, in particular, which is likely to be explained by the varying L2 proficiency, which correlated with the N400 amplitude and therefore may explain the observable variance.

##### N400: Time window between 300-450ms

ERP amplitude at the Central pool in the time window between 300-450ms was predicted significantly by language switch, (*B* = −0.44, 95% CI [−0.73, −0.15], *t*(27) = −2.95), even after controlling for age (*B* = 0.26, 95% CI [−0.32, 0.84], *t*(27) = 0.88). This means that when reading sentences that contained a language switch from L2 to L1 – independent of congruency –, the N400 effect was larger; that is, values were more negative than those for no-switching. There was no significant effect of congruence (*B* = −0.27, 95% CI [−0.56, 0.03], *t*(27) = 1.81). A linear regression additionally showed that the magnitude of the N400 – computed as no-switch minus switch amplitude (N400 effect becomes positive) – correlated with L2 proficiency (*B* = −0.44, 95% CI [−0.89, 0.00], *t*(18) = −2.10). Thus, for learners with lower L2 skills after the course, the N400 effect was larger than for learners with high L2 skills (see Figure 6). There was no significant effect of any of the factors included in the mixed model on N400 latency.

**Fig. 6.**
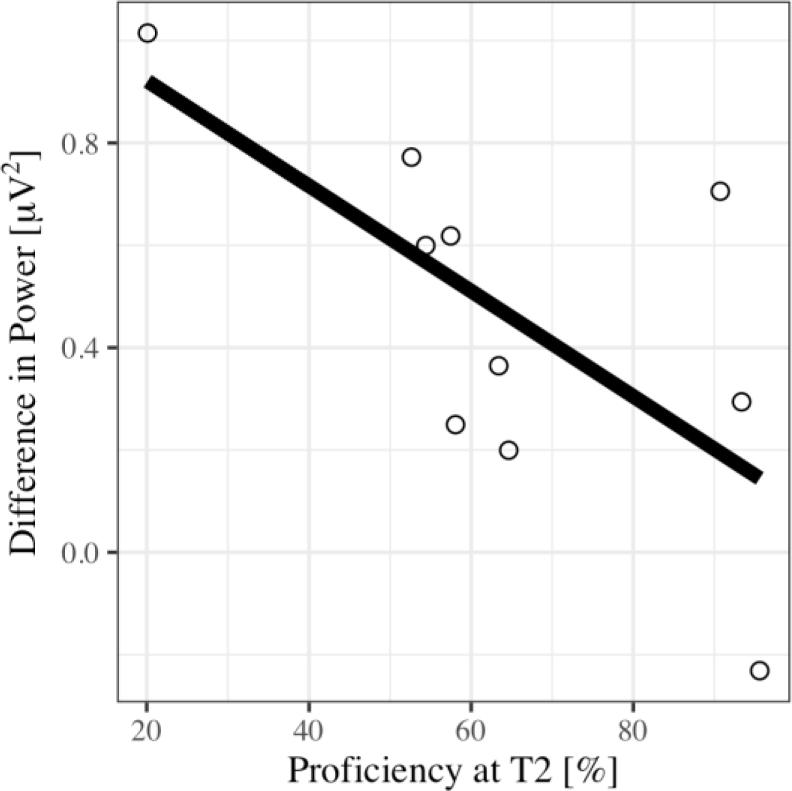
Regression plot for L2 proficiency at T2 and the N400 effect calculated as the difference between no language switch and language switch conditions, such that positive values on the y-axis represent the size of the fact (negativity inverted).

##### LPC: Time window between 600-800ms

The multilevel model for the ERP time window between 600-800ms at the Parietal pool showed no significant effect of age (*B* = 0.00, 95% CI [−0.60, 0.59], *t*(27) = 0.00) or language switch (*B* = 0.22, 95% CI [−0.11, 0.55], *t*(27) = 1.34), but a significant effect of congruence (*B* = 0.57, 95% CI [0.25, 0.90], *t*(27) = 3.40). This effect reflects the fact that incongruent conditions, whether in L1 or L2, elicited a larger LPC than congruent conditions. The correlation between LPC magnitude and proficiency at L2 was non-significant (*B* = −0.12, 95% CI [−0.61, 0.37], *t*(18) = −0.61). None of the factors in the mixed model had a significant effect on latencies of the LPC.

### 3.3. Discussion of Experiment 2

This experiment investigated behavioral and electrophysiological markers of L2 learning in old adulthood by examining the event-related potentials from a group of (Swiss-) German older learners (65-74 yrs) who participated in an intensive 3-week EFL training. We aimed to assess ERP correlates of L2 proficiency in order to study the relationship between L1 and L2 in older EFL beginners. As predicted from Van der Meij et al. (2011), language switching (from L2 to L1) elicited an N400 effect in the time window between 300-450ms. However, switching from L2 to L1 in our data did not elicit a LPC as it did in Van der Meij et al. (2011), whereas the additional condition of semantic incongruence – which was not present in the study by Van der Meij et al. (2011) – did.

In the 300-450ms time window, language switching (from L2 to L1) as compared to no-switching yielded an N400 effect over the Central electrode pool, independent of semantic congruency. The time window, the observed effect as well as its estimated location on the scalp are consistent with the findings of Van der Meij et al. (2011). Given that all our participants were L2 beginners, however, there was nothing to be gained by splitting the group into a low- and a high-proficiency group, as was done by Van der Meij et al. (2011). Nevertheless, the observed N400 effect could be shown to correlate with L2 proficiency after the training in that learners with a higher proficiency showed smaller N400 effects.

The N400 component, which originally was believed to be a marker of semantic deviation (Kutas & Hillyard 1980) and word frequency (Barber, Vergara, Carreiras 2004), has meanwhile also been shown to be elicited by language switching from L1 to L2 (Proverbio, Leoni & Zani 2004). The present study not only confirms that language switching back into the L1 can also yield an N400 effect, but that this effect even occurs in older learners, and does so after a training period of only three weeks. Given the low proficiency of our learners, the observed effect cannot be explained by frequency effects, because words in the L1 are conspicuously more frequent in the learners’ language experience than the recently learned L2. Instead, the N400 appears to be a marker of how active the two languages are at once and how much activation cost is required to switch from one to the other. As predicted, in our data, the N400 negatively correlated with L2 proficiency at T2, so that more advanced learners (i.e. lower intermediate level) showed smaller amplitude differences than learners with very basic L2 skills. The fact that the N400 as a function of language switch was reduced in the more proficient learners of our sample may be an indication that for them, the coactivation of L1 and L2 is stronger than in the less proficient learners. This finding is in line with the effects observed in Van der Meij et al.’s intermediate learners (2011). At the same time, however, Van der Meij et al. (2011) also found that switching costs increased again for very advanced L2 speakers. Thus, it appears that L2 and L1 lexicons are strongest coactivated at intermediate stages of L2 learning. At initial stages, the L2 lexicon does not yet exist, while at a very high proficiency, the L1 is inhibited for successful L2 processing (Abutalebi 2008; Cuppini et al. 2013), both of which explain the lexical surprise effect, as typified by the N400. In contrast to Van der Meij et al. (2011), we argue that these effects can be explained without assuming separate lexicons for L1 and L2, or selective language access, as postulated in the Revised Hierarchical Model (Kroll & Stewart, 1994). This model has repeatedly come under attack (Kroll, Van Hell, Tokowicz, & Green, 2010; Brysbaert & Duyck, 2010), and it is likely that the L1-L2 coactivation in our intermediate learners does not point to a link between L1 and L2 lexicons only, but that it manifests as an indirect association of both languages with the conceptual system through episodic memory, in particular because the interference of L1 words on L2 processing has been shown to be modifiable through the global language context, and is, thus, far from stable (Elston-Güttler, Gunter, & Kotz, 2005).

As mentioned above, however, differences between the participants in our study and those in the study of Van der Meij et al. (2011) were not limited to L2 proficiency, but also applied to the participants’ ages, and this difference was reflected in the ERP data. Compared to the results of Van der Meij et al. (2011), who performed a similar experiment on language switching in younger adults, we found that ERP amplitudes in general were noticeably smaller in our group of older adults. This finding is consistent with the results of Xu et al. (2017), who found that the congruency-induced N400 (in L1) yielded significantly smaller N400 amplitudes for older compared to younger adults.

One reason for this reduction in ERP amplitudes could be the age-related brain atrophy (Giroud et al., 2019), however, it is not yet clear how these differences in amplitude may relate to behavioral losses. Another explanation could be that greater variability in the peak-evoked amplitudes resulted in smaller averaged ERPs, or that the reduction in ERP amplitude reflects a shift in how the stimuli are processed.

A further difference between our data and those of Van der Meij et al. (2011) is that we did not observe a significant LPC in response to language switching. In this case, the absence of a switching-related LPC could be due to the age difference or the difference in L2 proficiency between our study and that of Van der Meij et al. (2011). A recent study by Kim, Oines and Miyake (2018) would speak in favor of the former, as they found that verbal working memory capacities correlated positively with LPC amplitudes and negatively with N400 amplitudes, both being generated by the same semantic anomalies and occurring within the same individuals but in different markedness. Since working memory capacities commonly affected by the age-related cognitive decline (e.g. Salthouse, 2010), reduced working memory could likely be responsible for the absence of a switch-related LPC in our sample.

We did, however, observe a late positivity with a parietal distribution that varied as a function of semantic incongruence, a condition that we added to the study design of Van der Meij et al. (2011). Considering the high accuracy in the comprehension questions, the observed LPC indicates that learners processed the stimulus sentences for content, too, not only word by word. Here, we employ the term LPC for reasons of terminological consistency with Van der Meij et al. (2011). However, as noted repeatedly in the P600 literature, the terms “(semantic) P600”, “late-positive shift”, “late-positive effect” or “late-positive component” can be used interchangeably, as it is likely that they are attributable to the same underlying neurobiological processing mechanisms, independent of a preceding N400 effect (Bornkessel-Schlesewsky et al. 2011; Dröge, Fleischer, Schlesewsky, & Bornkessel-Schlesewsky 2016). The latency of the incongruence-effect as late as 600-800ms can be explained by the fact that in order to detect the semantic incongruence, lexical processing of the critical word needed to be completed, a computation that is performed only in the N400 time window. In line with this, visual inspection of the grand average ERP indicated that semantic incongruence also modulated the N400 in the no-switching conditions, suggesting that semantic processing took place earlier when no switching was required. This effect, however did not reach significance, which in turn would be in line with studies showing a reduction of N400 effects with age (Gunter, Jackson & Mulder, 1998). It is therefore not entirely clear whether we are dealing with a biphasic N400 - late positivity pattern or a monophasic LPC.

## 4. Overall Discussion and Conclusion

The field of second-language acquisition in old adulthood is still in its infancy, and consequently there is little to no research on the neurophysiological markers of L2 learning in old age. In this pilot study, we have been able to replicate two studies on younger adult L2 learners with a sample of older (Swiss-) German learners of English as a foreign language. Our resting-state data replicated the study by Prat et al. (2016), confirming the role of the beta1 band in L2 learning, transferring the same research design from younger to older L2 learners (see also Küssner et al., 2016). We could show that beta1 power before and during the training predicted the L2 development. These findings feed into the discussion around L2 learning aptitude, suggesting beta1 power as a possible electrophysiological correlate thereof. Since beta1 power has been associated with selective attention and semantic working memory, our findings are consistent with psycholinguistic theories on L2 learning (Collins, Trofimovich, White, Cardoso, & Horst, 2009; Ellis, 2006). In agreement with these theories, successful learning at initial stages of the L2 training may depend on beta1 oscillations that correspond to selective attention and semantic working memory in order to extract and focus on novel forms and meanings from the L2 input. Possible future applications of these findings are manifold. For instance, it remains for future research to show whether the L2 development can be increased by enhancing beta1 oscillations before each learning session through neurofeedback, whether the training can be adjusted to individual differences in beta1 power or whether beta1 oscillations themselves can be enhanced through the L2 training.

Second, we assimilated the research design by Van der Meij et al. (2011), and could confirm that, even after a short training period of only three weeks, older L2 learners also manifested an N400 effect towards language switching from the newly learned L2 into their L1. Language switching appeared to require less effort with increased proficiency, suggesting that the lexicons of L1 and L2 had already started to become more closely linked, which confirms the theories of Abutalebi (2008) and Cuppini et al. (2013). According to those theories, the L2 parasitizes its L1 equivalent at initial stages of L2 learning, and the two systems only become independent language systems again with highly advanced proficiency, thus indicating that integration processes of L1 and L2 change with the individual competence in the L2.

To sum up, our findings confirm that the human language system remains malleable even until third age and appears to follow the same electrophysiological mechanisms we observe in younger adults. The question that we plan to address in the future is not only how good older adults are at learning new languages, but also whether and to what extent language learning can be beneficial for an individual third age person (see Pfenninger and Singleton, 2019, for state-of-the-art article).

## 5. Limitations

The data presented in the present study are not without limitations. Even though our findings constitute an important first step towards understanding the neurophysiological mechanisms underlying L2 learning in old adulthood, the present design only reports on differences between pre- and post-training data, largely ignoring the learning process occurring between the two. Thus, in order to understand the individual L2 learning trajectories, dense longitudinal studies throughout longer periods are required that allow inferences, for instance, as to whether beta1 power is more predictive of L2 development at specific stages of the learning process. Accordingly, here we only report findings from a group of older L2 beginners, and even though these findings are theoretically well compatible with those of Van der Meij et al. (2011), future studies will have to show whether the differences between our findings pertain solely to differences in L2 proficiency, or alternatively to differences in age, training type, sample size, language tests etc. We did not include a younger control group in the present study because our focus was on individual differences *within* the population of older learners themselves. We judged that differences between this group and a younger control group would be of little informational worth, as it would be impossible to answer whether such differences occurred due to the degenerative processes of aging itself or to experientially determined differences in neurobiology and cognition. It has repeatedly been shown that although on average, cognitive performance declines with age, the variability of this construct is so great amongst older adults, that the cognitive functioning of some remains comparable to that of younger adults (Whalley et al., 2004). Accordingly, here we focused on individual variability in the ability to acquire a new language in third age and do not make any claims as to whether or not the observed effects are specific to language learning itself. We therefore also refrained from including an age-matched control group.

Moreover, in this study we used one single measure of L2 proficiency to assess general L2 skills, and will investigate in future studies how our findings translate into different language domains (speaking, listening, writing, reading), and whether, for instance, beta1 power is predictive of L2 development in each of them.

Finally, the sample size in this pilot study was constrained so that the instructor could attend to each of the learners’ questions and needs, especially because the extent of interindividual differences was unclear beforehand. Therefore, We interpret our results with caution and plan to replicate them in a more comprehensive study that is currently being conducted in our lab.

## Acknowledgements

We would like to express our gratitude to Ms Allison Christen for her invaluable comments on earlier version of this manuscript, and to the URPP “Dynamics of Healthy Aging” for providing the required infrastructure throughout all phases of the project.

## Data Availability

The data that support the findings of this study are available from the corresponding author, MK, upon reasonable request.

## Abbreviations

CEFR: Common European Frame of Reference
CorP: Corrected Progress
EEG: Electroencephalogram
EFL: English-as-a-foreign-language
ERP: Event-related potentials
L1: Native language
L2: Foreign language
LPC: Late-positive component
T1: Time point one (pre-training)
T2: Time point two (post-training)

